# Analysis of menstrual effluent uncovers endometriosis-specific cell populations and impaired cellular pathway processes

**DOI:** 10.1101/2025.08.21.671582

**Authors:** Taylor R. Wilson, Stephanie A. Morris, Paul L. Deford, Andrea Starostanko, Susan Kasper, Simran Venkatraman, Caroline F. Morrison, Kathryn Friend, Seddon Y. Thomas, Katherine A. Burns

**Affiliations:** Division of Environmental Genetics and Molecular Toxicology, Department of Environmental and Public Health Sciences, University of Cincinnati, Cincinnati, OH; Reproductive Endocrinology and Infertility, Department of Obstetrics and Gynecology, University of Cincinnati, Cincinnati, OH; College of Nursing, University of Cincinnati, Cincinnati, OH; Thermo Fisher Scientific, Durham, NC

## Abstract

Endometriosis is a chronic gynecological disease affecting 1 in 10 reproductive-aged women and is characterized by the ectopic presence of endometrial tissue outside the uterus. The leading hypothesis for disease etiology is via the reflux of menstrual effluent (ME) into the peritoneal cavity. ME is a complex mixture of viable endometrial tissue, proteins, and immune cells which serve specialized functions during menstruation to support and repair the endometrium. We hypothesized shifts in the subpopulations of immune cells present during retrograde menstruation may alter the microenvironment of the peritoneal cavity leading to a favorable environment for uterine tissue attachment and survival of endometriotic lesions. Menstrual effluent collected on days 1 and 2 of menstruation identified from women with endometriosis distinct morphological features, increased subpopulations of aged neutrophils, increased anti-inflammatory macrophages, and overall impaired clearance pathways that hijack endometrial clearance likely contributing to the development and progression of endometriosis.

## Introduction

Endometriosis is a chronic gynecological disease affecting 1 in 10 reproductive-aged women worldwide (1, 2). Symptoms include dysmenorrhea, dyspareunia, excessive bloating, non-menstrual pelvic pain, digestive problems, infertility, and increased risk of autoimmune disorders (3–6). Endometriosis is characterized by growth of endometrial tissue outside of the uterus which contains glands, stroma, fibrotic tissue, and hemosiderin ladened macrophages that form ectopic lesions (1, 7). The current diagnostic for definitive confirmation of endometriosis requires invasive laparoscopic surgery for histopathological examination of the lesions. The difficulty and cost required in obtaining a diagnosis results in a diagnostic delay of 7-9 years (8).

While the exact etiology of endometriosis is unknown, a multifactorial paradigm encompassing environmental, genetic, hormonal, and immunological factors is likely (9–11). The leading hypothesis for disease etiology is via retrograde menstruation where menstrual effluent (ME) flows retrograde, back though the oviducts and into the peritoneal cavity (12). While nearly all women experience retrograde menstruation, only 10% develop endometriosis; thus, aberrant regulation of the immune system may alter conditions of the peritoneal microenvironment to accommodate ectopic lesions (1, 2, 9, 13). Although endometriosis is not identified as an autoimmune disorder, women with endometriosis have increased risks for autoimmune diseases and immune dysfunctions are observed systemically and within the peritoneal cavity (14–18). Literature supports defects in the immune system may facilitate exacerbation or early initiation stages of endometriosis lesion formation from changes in immune-cell recruitment, cell-migration, cell-adhesion, angiogenesis, and inflammatory processes (11).

Endometriosis studies widely utilized peritoneal fluid, peripheral blood, endometrial biopsies, and surgically excised lesion tissues to examine the white blood cell (WBC) populations and functions associated with disease pathology (19–22). Of note, these methods of sample collection are invasive and often cause discomfort or pain during collection. ME is a less invasive method of sample collection and may be instrumental in understanding the functional role of the immune system during antegrade and retrograde menstruation. Menstruation is a highly inflammatory process that consists of structural changes occurring in the uterine microenvironment to endometrial composition and the recruitment of WBC populations (23). ME is composed of endometrial, red blood, endothelial, and WBCs—with 40% of WBCs being neutrophils, macrophages, and uterine natural killer cells (13, 24).

The menstrual cycle has three main phases and with each phase the uterine lining and the recruitment of WBCs changes. An influx of WBCs primes the uterine microenvironment prior to either implantation or the onset of menstruation (23, 25). These populations and associated cytokines are present in antegrade menstruation and are a representation of the composition of retrograde menstruation (23, 24, 26). While prior studies characterized and/or quantitated WBC populations in the uterus, little is known regarding their intricate actions during menstruation or their potential altered functions in gynecological disease (2, 23). During a normal menstrual cycle, various WBC populations scavenge for endometrial tissue debris and remodel the endometrial lining (23, 25). In contrast, a disruption to the composition of the WBC populations would skew the inflammatory response leading to dysfunction(s) within the uterine and peritoneal cavity microenvironments. In endometriosis patients, immune deficits propose defective clearance of endometrial debris by phagocytes, reduced functional activity in WBCs, and inflammation within the peritoneal cavity (2, 23, 27, 28). Together, these studies underscore the likelihood of immune-driven factors to contribute to the complex disease pathology of endometriosis.

Endometriosis patients have an increased risk for comorbidities of autoimmune diseases attributed to dysfunctions in the innate and adaptive immune system (28–31), suggesting altered immunological responses are involved in the pathogenesis of endometriosis or endometriosis lesions exacerbate the local inflammatory environment causing dysfunction. For example, neutrophils and neutrophil-associated cytokines increased 3 to 5-fold in the peritoneal fluid of women with endometriosis (11, 28, 32, 33); however, few studies examined neutrophils and other WBCs in ME (1, 24, 26, 34). Together, known immunological dysfunctions occur in women with endometriosis and these WBC subpopulations will be important for delineating the inflammatory responses which may exacerbate endometriosis.

Since ME is a monthly occurrence and can be collected with ease and limited discomfort, it was chosen as a substrate and as a diagnostic tool. ME is distinct from peripheral blood, but is, consistent with endometrial biopsy specimens with a higher yield of WBCs (26, 35). We hypothesized that ME from women with endometriosis compared to healthy controls would exhibit a unique immune profile predominated by changes in neutrophil and macrophage subpopulations. Thus, we characterized WBCs in ME by examining subpopulations through deep immunophenotyping and differential staining, and examined the acellular fraction of ME for cell free DNA (cfDNA) and proteomic analysis. Our data provide evidence the ME from women with endometriosis compared to healthy controls have increased populations of aged neutrophils, anti-inflammatory macrophages, and T helper cells whereas control ME has elevated proinflammatory macrophages and cytotoxic T cells. Neutrophils likely act as effector cells through premature aging and increased angiogenic and adhesive cell surface markers leading to a possible dysfunctional menstrual microenvironment, thus, likely shifting the inflammatory response in the peritoneal cavity during retrograde menstruation contributing to the initiation and progression of endometriotic lesions.

## Results

### Menstrual effluent participant demographics and menstrual history

Participants were asked to provide ME on days 1 and 2 of menstruation and complete a health questionnaire designed using the World Endometriosis Research Foundation (36) guidelines (Table 1). Study participants were separated into groups based on condition (healthy control, endometriosis, and endometriosis with exogenous hormonal therapies). The participants ranged from 21 to 44 years old with no statistical difference between groups. Weight and height were used to calculate BMI and no differences were observed among the groups. Gravidity and parity, including the number of pregnancies, live births, and miscarriages, were not different among groups. The primary population of participants were Caucasian/white and is a limiting factor in the diversity of this study. Although gravidity and parity did not differ, women with endometriosis had decreased fertility. Endometriosis women had complications with infertility for more than 1.5 years before pregnancy whereas the participants in the control group with infertility experienced infertility complications lasting less than 1.5 years. No difference was observed between menstrual cycle length and duration of flow length. Nearly every participant in the endometriosis groups had a history of pelvic pain compared to 10% of control participants. Interestingly, the range of symptoms was variable between individuals ranging from no symptoms to all symptoms asked. Menstrual cramping, digestive problems, excessive bloating, lower back and/or leg pain, and dyspareunia were significantly increased in women with endometriosis compared to the healthy control group. Non-menstrual cramping, nausea and/or vomiting, diarrhea and/or constipation, painful bowel movements, severe migraines, and sudden mood swings were not different among conditions. While not significant uncontrollable crying (p = 0.0769) trended to be higher in women with endometriosis. Overall, these data support previously published studies on endometriosis that differences exist between control and endometriosis groups suggesting that menstruation for women with endometriosis, regardless of hormonal therapies, has greater symptomatic severity and increased infertility.

**Table 1.**
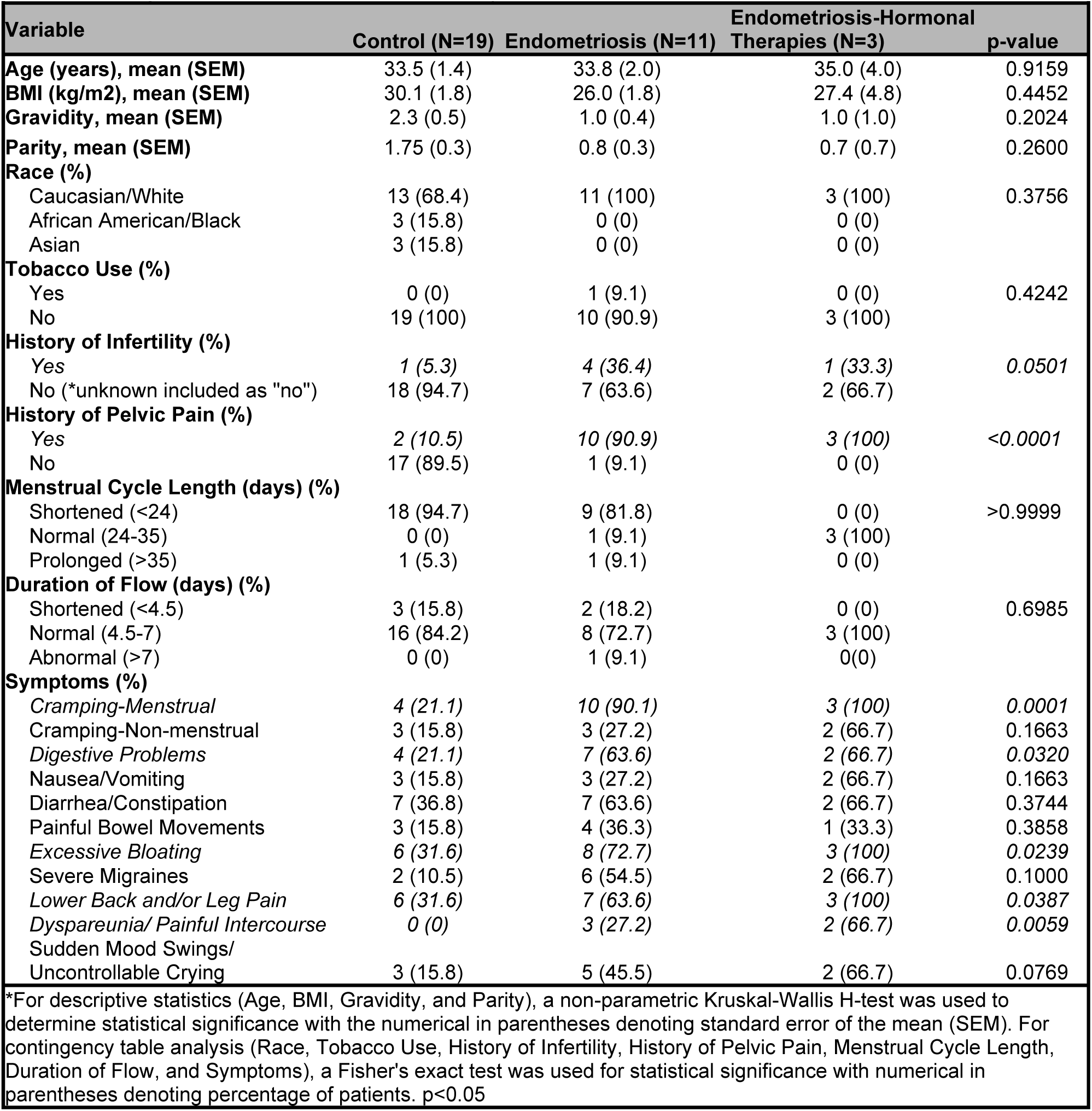
Demographics of Menstrual Effluent Study Participants.

### Menstrual flow rates are higher in women with endometriosis

ME samples were collected on day 1 and 2 of menstruation with collection beginning at full flow once spotting finished. Sample volumes varied between individuals (Figure S1A) with no difference in the amount of ME collected among condition or day. Each participant was asked for an estimated time of sample collection to account for sample collection and individual variability. Sample collection times ranged from 6-24 hours per day. To normalize sample collection and determine the menstrual flow rate, the amount of ME collected (mL) per day was divided per time (hour) of sample collection for each individual and day (Figure S1B). In each condition, day 1 was not significantly different to the paired day 2 of the same condition. However, for endometriosis participants the volume on day 1 and 2 was approximately 1.5-fold higher than control day 1, control 2, and endometriosis with exogenous hormones day 2 and approximately 2-fold higher than endometriosis day 1. The endometriosis with exogenous hormones group was expected to have a lower flow rate due to the influence of hormonal therapies on endometrial thickness (37). To determine if day 1 flow rate was different than day 2 per individual, the ratio of day 1 divided by day 2 was calculated (Figure S1C). Individually, the ratio of day 1 to day 2 flow rates were not different from one another, indicating that individuals have similar flow rates on days 1 and 2. Overall, the increased flow rate during menstruation may contribute to the severity of symptoms in women with endometriosis.

### A morphologically distinct neutrophil phenotype is observed in ME of women with endometriosis

As our hypothesis was to focus on the WBCs in ME, only WBCs were isolated from ME for deep immunophenotyping. An aliquot of WBCs from each day was used for cytological visualization. The cytology demonstrates the full complement of WBCs are present in ME (i.e., neutrophils, macrophages, eosinophils, and B/T cells) (Figure 1). Like peripheral blood, neutrophils are the most abundant WBC type present in the ME. Neutrophils have a distinct cellular morphology of a polymorphonuclear or multi-lobed nucleus and stain dark purple in the nucleus and a light pink to violet in the cytoplasm. In control day 1 and 2 images (Figure 1A-B), neutrophils exhibit 2 to 3 nuclear lobules with few to no vacuoles in the nucleus or cytoplasm. In stark contrast, in the endometriosis samples (Figure 1C-D), the neutrophils are often hypersegmented and have greater than 3 lobules with as many as 6 to 8 per cell. The hypersegmented phenotype in neutrophils depicts a morphologically aged maturation state of the cells (38, 39). Additionally, the neutrophils in women with endometriosis have small clear vacuoles in their nuclei and cytoplasm which additionally suggests an altered physiological or activation state. Critically, the use of exogenous hormonal therapies did not alter the aged phenotype or appearance of the neutrophils in women with endometriosis (Figure 1E-F) as their morphology is similar to the morphological features observed in endometriosis participants not on exogenous hormones. The distinct appearance of the neutrophils in women with endometriosis compared to healthy control women demonstrates that a morphological phenotype may serve as a noninvasive method to diagnose endometriosis and that a physiological difference may underlie a functional defect and/or altered inflammatory state in women with endometriosis leading to the increased likelihood of lesion development.

**Figure 1.**
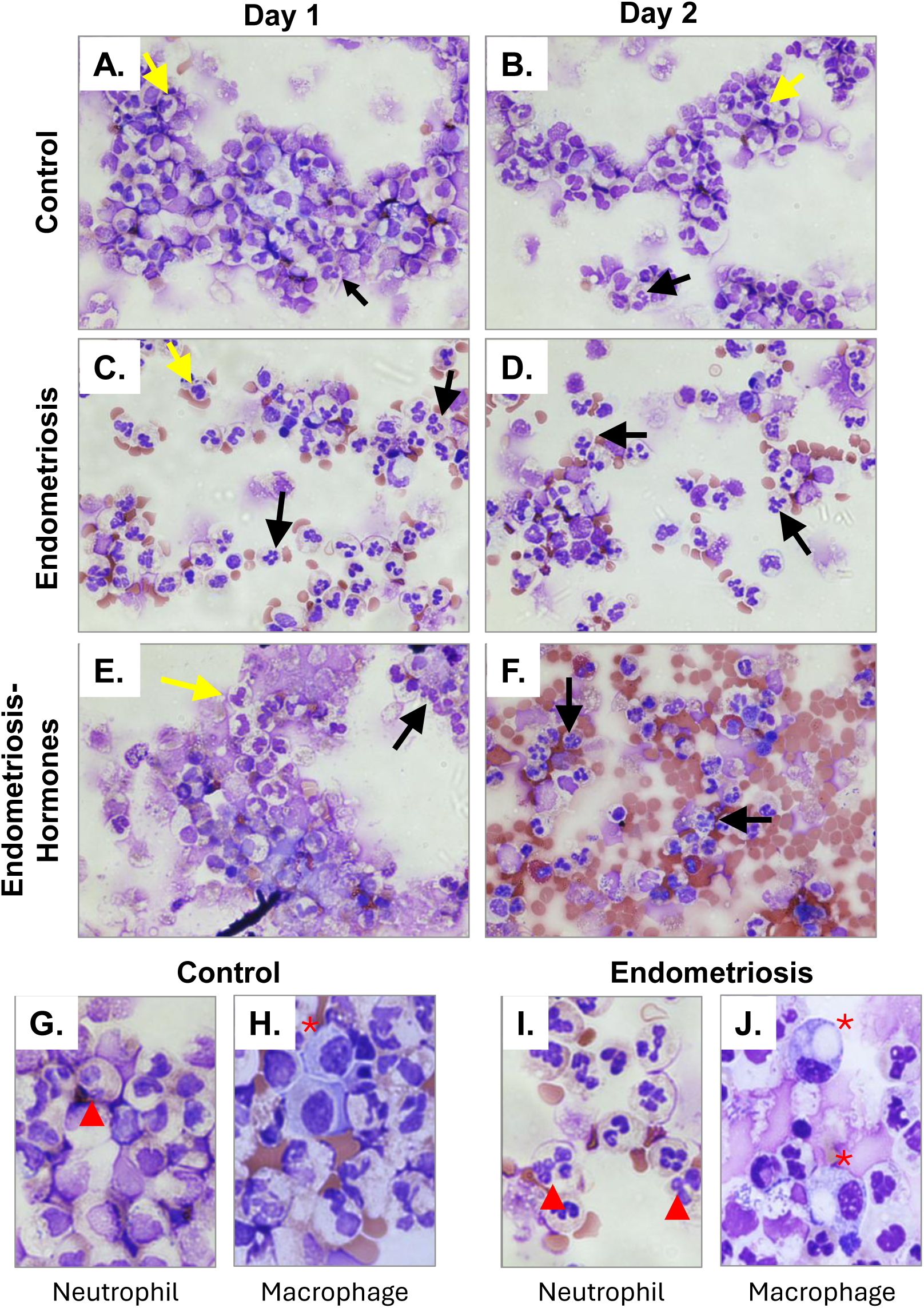
Menstrual effluent from women with endometriosis have morphologically distinct hypersegmented neutrophils with vacuole structures. Differentials of isolated WBCs from ME were stained with modified Giemsa (Hema III). **(A-B)** Control days 1 and 2. Neutrophils (yellow arrows) have 2-3 nuclear lobes with few to no vacuoles. Occasionally, hypersegmented neutrophils (4+ lobes, black arrow) and vacuoles are seen in control women. **(C-D)** Endometriosis days 1 and 2. Neutrophil (black arrows) nuclei are hypersegmented and contain vacuoles (to the top of red arrowhead). **(E-F)** Endometriosis with exogenous hormonal therapies days 1 and 2. Neutrophils (black arrows) nuclei are hypersegmented and contain vacuoles. **(G** and **I)** Inserts of **A** Control and **C** Endometeriosis to denote vacuoles (red arrowhead). **(H** and **J)** Inserts of macrophages from control and endometriosis to denote foamy macrophages. Control (n=19); Endometriosis (n=11); Endometriosis-Hormones = endometriosis with exogenous hormones (n=3).

In addition to the altered neutrophil phenotype, albeit in fewer numbers, is a change in the macrophage phenotype. The predominant morphology of the macrophages found in ME from women with endometriosis have an inflated and highly activated appearance with sizeable vacuoles within the cytoplasm (Figure 1 H and J). This macrophage phenotype is often denoted as “foamy macrophages”. Foamy macrophages are typically associated with atherosclerosis; however, they are linked to anti-inflammatory functions to suppress inflammation in inflammatory conditions and accumulate lipids within vacuoles to create the foamy appearance (40–42). This shift in phenotype in ME from women with endometriosis compared to control women indicates a potential compensation to mitigate the altered inflammation during menstruation.

### Granulocyte subphenotyping reveals differential recruitment of neutrophils in women with endometriosis

We showed previously, aged and angiogenic neutrophil populations increased in ME on day 1in endometriosis compared to day 1 of healthy controls (43). This study expands on the previously identified population by further characterizing neutrophil subtypes (Figure S2) and includes a second day of collection to identify shifts in WBC populations.

We first examined total neutrophils on days 1 and 2 of menstruation and compared healthy control participants, endometriosis participants, and endometriosis participants on exogenous hormones. The total neutrophil ratio is 85-95% of the total WBC population in ME (Figure 2A). The total neutrophil ratio did not differ when comparing condition or day. Shifts in external surface markers occur because of neutrophil plasticity; therefore, multiple external cell surface markers were used to capture subsets of these dynamic populations. Neutrophils were gated on CD16, CXCR2, and CXCR4 to characterize their levels of maturity on days 1 and 2 of menstruation. Mature neutrophils express CXCR2 and as they transition to an aged phenotype, they will gradually express CXCR4. Mature neutrophils did not differ in quantity between days 1 and 2 of control; however, mature neutrophils increased recruitment on day 2 compared to day 1 in endometriosis and endometriosis with exogenous hormones (Figure 2B). Small shifts in subpopulations can be biologically significant, thus, a 19% increase in mature neutrophils on day 2 in endometriosis may hold biological significance.

**Figure 2.**
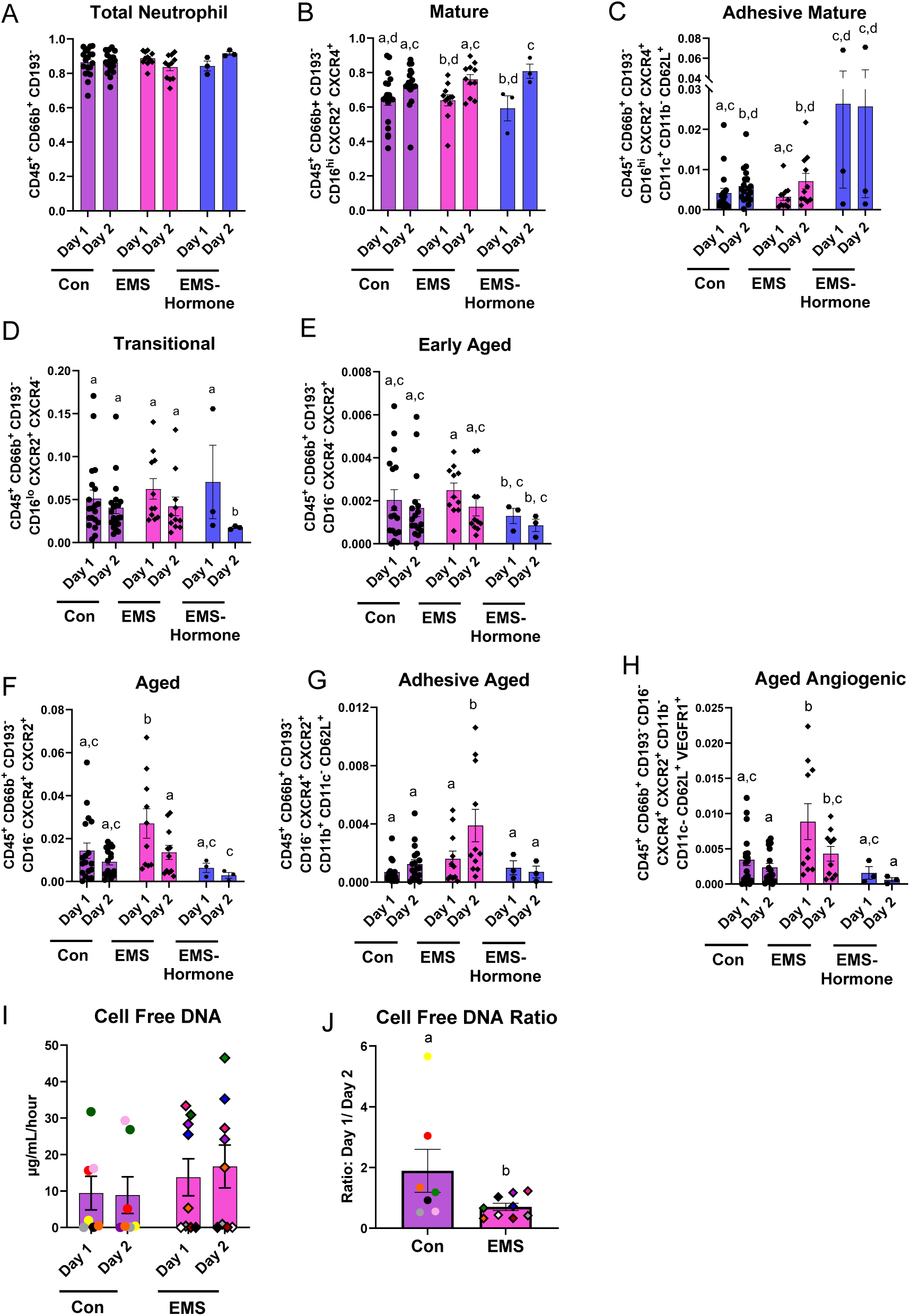
Women with endometriosis have higher levels of aged neutrophil subpopulations compared to control women. **(A)** Quantitation of total neutrophils to total CD45^+^ ratio. Total neutrophils were defined by: CD45^+^ CD66b^+^ CD193^-^. **(B)** Subpopulation ratios were calculated to total neutrophils. Mature neutrophils were defined as: CD45^+^ CD66b^+^ CD193^-^ CD16^hi^ CXCR2^+^ CXCR4^+^. **(C)** Adhesive mature neutrophils were defined as: CD45^+^ CD66b^+^ CD193^-^ CD16^hi^ CXCR2^+^ CXCR4^+^ CD11c^+^ CD11b^-^ CD62L^+^. **(D)** Transitioning neutrophils were defined as: CD45^+^ CD66b^+^ CD193^-^ CD16^lo^ CXCR2^+^ CXCR4^-^. **(E)** Early aged neutrophils were defined as: CD45^+^ CD66b^+^ CD193^-^ CD16^-^ CXCR4^-^ CXCR2^+^. **(F)** Aged neutrophils were defined as: CD45^+^ CD66b^+^ CD193^-^ CD16^-^ CXCR4^+^ CXCR2^+^. **(G)** Adhesive aged neutrophils were defined as: CD45^+^ CD66b^+^ CD193^-^ CD16^-^ CXCR4^+^ CXCR2^+^ CD11b^+^ CD11c^-^ CD62L^+^. **(H)** Aged angiogenic neutrophils were defined as: CD45^+^ CD66b^+^ CD193^-^ CD16^-^ CXCR4^+^ CXCR2^+^ CD11b^-^ CD11c^-^ CD62L^+^ VEGFR1^+^. Control (*n* = 19); endometriosis participants (*n* = 11); endometriosis with hormones (*n*=3). Data represent ± SEM. Statistical significance for each graph was determined by two-way ANOVA, followed by nonparametric, Mann-Whitney, 1-tailed *U* test. ^#^*p*<0.05. **(I)** Cell free isolated from the acellular fraction of ME from day 1 and 2 of sample collection. **(J)** Ratio of cell free DNA (day 1/ day 2). Control (n=19), Endometriosis (n=11), Endometriosis with Hormones (n=3). Data represent ± SEM. Statistical significance for each graph was determined by nonparametric, Kruskal-Wallis test followed by Mann-Whitney, 1-tailed U test. *p*<0.05.

Next, the mature neutrophils were examined for CD62L (a marker for recruitment and adhesion) (44) and denoted as adhesive mature neutrophils (Figure 2C). This population increased from day 1 to day 2 in both the control and endometriosis groups, demonstrating this population is functioning and/or is recruited similarly in healthy controls and women with endometriosis. In contrast, this subpopulation was variable in endometriosis with exogenous hormones and, although the data suggest significance, the small sample size likely skews how this population is perceived. These findings suggest the adhesive mature neutrophil subpopulation may differ in their recruitment from exogenous hormones.

The transitional maturity state of neutrophils was defined by lower levels of CD16 (CD16^lo^) and lack of CXCR4 (CXCR4^-^) as this population is shifting towards a more aged phenotype (Figure 2D). This population did not differ between day or condition, except for day 2 in the endometriosis group with exogenous hormones which had a 50% decrease compared to control day 2.

The differentials display a morphological difference in the number of lobules and vacuoles present in endometriosis samples. Based on these observations, neutrophils in women with endometriosis were suspected to be aged. The activation states of neutrophils change due to their inflammatory environment; therefore, capturing every aged subphenotype of neutrophils would be challenging. However, collectively these aged subpopulations of neutrophils may morphologically represent the distinct neutrophil phenotypic appearance found in Figure 1. The early aged population was based on the absence of CXCR4 (Figure 2E). This population did not differ between the control and endometriosis groups; however, endometriosis day 1 was 2- and 3-fold higher than endometriosis with exogenous hormones on days 1 and 2, respectively.

A more classically aged immunophenotype of neutrophils was identified by the absence of CD16 and the presence of both CXCR2 and CXCR4 (Figure 2F). The more classically aged neutrophils do not differ from day 1 to day 2 in healthy women; however, in women with endometriosis, day 1 increased compared to control days 1 and 2, endometriosis day 2, and both endometriosis with exogenous hormones. Endometriosis day 1 increased by 2-fold compared to control day 1 and endometriosis day 2 as well as nearly 3-fold compared to endometriosis day 2 and endometriosis with exogenous hormones on day 1. The endometriosis with exogenous hormones on day 2 is significantly lower than both endometriosis days which suggesting exogenous hormones may impact this subpopulation. The classically aged population was further characterized by the presence or absence of an integrin (CD11b) and a selectin (CD62L) to be markers of an adhesive phenotype. The population (Figure 2G) does not differ from day 1 to day 2 in healthy control women; however, in women with endometriosis, an increase in recruitment is observed on day 2 compared with each other condition and day with nearly 2-fold the abundance compared to control day 2. These findings indicate a shift in the inflammatory profile in day 2 of endometriosis may promote and/or increase adhesive properties of ME that flows antegrade and retrograde. While aged angiogenic neutrophils were identified previously (43), we further characterized this subpopulation by the absence of CD11b and the presence of CD62L and VEGFR1 (Figure 2H). Within the three groups no significant shift occurred between days 1 and 2 of menstruation; however, endometriosis day 1 had 2- and 4-fold greater recruitment compared to control day 1 and endometriosis with hormones day 1, respectively. The levels of aged angiogenic neutrophils in endometriosis day 2 is ∼2- and 8-fold higher compared to control day 2 and endometriosis with hormones on day 2, respectively. This increase observed in the endometriosis group demonstrates factors are present in the ME that promote the externalization of angiogenic cell surface markers, but not in the endometriosis with exogenous hormones group suggesting that this aged angiogenic population may have physiological relevance in endometriosis pathogenesis. Collectively, the neutrophil subpopulations exhibit shifts in their external surface markers to suggest altered composition and/or signaling occurs in the ME of women with endometriosis.

### Cell free DNA release shifts in control from day 1 to 2 of menstruation

With senescence, apoptosis, and the release of extracellular traps from neutrophils, cfDNA is released into the extracellular space and cleared by phagocytic cells (45, 46). The acellular fraction of ME, separated before WBC isolation, was used for cfDNA isolation and proteomic analysis. Once cfDNA was normalized per person [amount (µg/mL) per hour collected], no differences were observed when comparing condition or day (Figure 2I). The cfDNA value per person was visually assigned a color as the range varied among individuals; a ratio was calculated to examine the quantity of change individually (Figure 2J). Control samples showed variability, but endometriosis had little indicating a disease specific change. The change in cfDNA levels from day 1 to 2 in controls is 2-fold the ratio of the endometriosis group, demonstrating the cfDNA released in the endometriosis group was constant. The increased shift in cfDNA released in controls from day 1 to day 2 suggests increased apoptosis or extrusion of NETs in control ME and lack of clearance in endometriosis as the ratio and levels do not change.

### Macrophage subphenotyping reveals distinct subpopulation recruitment in women with endometriosis

Macrophages are phagocytic cells which actively clear debris during menstruation, are largely monocyte derived, and are recruited by neutrophils (23, 25). Therefore, we next examined macrophages. Total macrophages account for about 10-15% of the WBC population (Figure 3A) and did not differ by condition or day of menstruation. Macrophages were further characterized by the presence (proinflammatory) or absence (anti-inflammatory) of CD16. Proinflammatory macrophage recruitment decreased in endometriosis day 2 by ∼40% compared to control days 1 and 2 (Figure 3B). The endometriosis with exogenous hormones group did not differ from control or endometriosis groups. The decrease in pro-inflammatory macrophages may be from recruitment or altered inflammatory responses due to neutrophil populations in ME on day 1 of endometriosis. Of note, a subpopulation of proinflammatory macrophages with the presence of integrins (CD11b and CD11c), a chemokine receptor (CX3CR1), and MHC class II (HLA-DR) was characterized as a pro-inflammatory antigen presenting macrophage (Figure 3C). Endometriosis day 1 and endometriosis with hormones day 1 had significantly less recruitment in comparison to control day 1 by 2-fold and 4.5-fold, respectively, with no difference between day 2 population abundance. These findings indicate a potential impairment in this particular proinflammatory phenotype in endometriosis may contribute to an altered immune responses or indicate that neutrophils or other factors play a larger role in the pro-inflammatory state on day 1.

**Figure 3.**
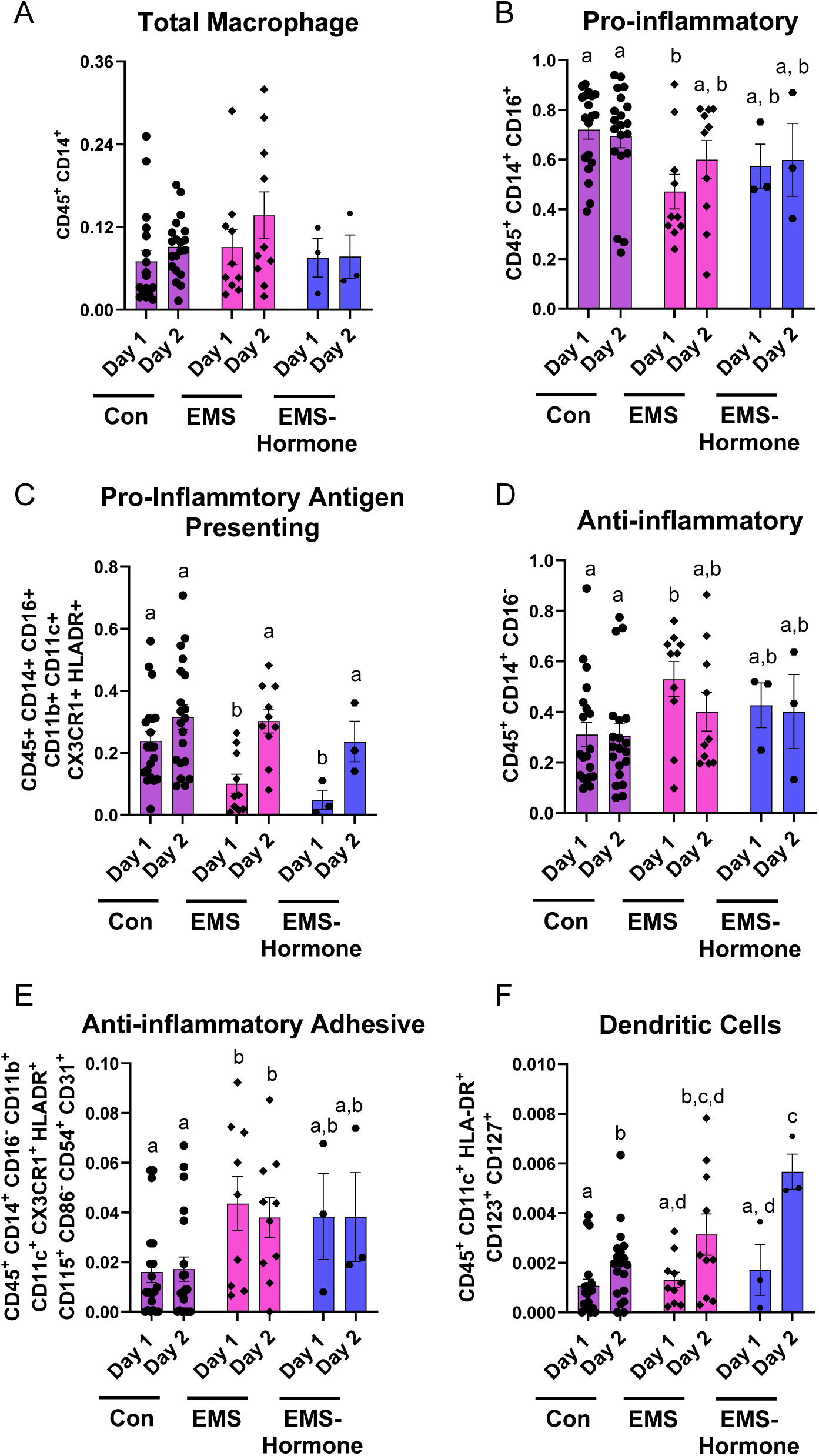
Women with endometriosis have increased anti-inflammatory macrophages. **(A)** Quantitation of total macrophages to total CD45^+^ ratio. Total macrophages were defined by: CD45^+^ CD14^+^. **(B)** Subpopulation ratios were calculated to total macrophages. Pro-inflammatory macrophages were defined as: CD45^+^ CD14^+^ CD16^+^. **(C)** Proinflammatory antigen presenting macrophages were defined as: CD45^+^ CD14^+^ CD16^+^ CD11b^+^ CD11c^+^ CX3CR1^+^ HLADR^+^. **(D)** Anti-inflammatory macrophages were defined as: CD45^+^ CD14^+^ CD16^-^. **(E)** Anti-inflammatory adhesive macrophages were defined as: CD45^+^ CD14^+^ CD16^-^ CD11b^+^ CD11c^+^ CX3CR1^+^ HLADR^+^ CD115^+^ CD86^-^ CD54^+^ CD31^+^. **(F)** Dendritic cells were defined as: CD45^+^ CD11c^+^ HLA-DR^+^ CD123^+^ CD127^+^. Control (*n*=19); endometriosis participants (*n* = 11); endometriosis with hormones (*n*=3). Data represent ± SEM. Statistical significance for each graph was determined by two-way ANOVA, followed by nonparametric, Mann-Whitney, 1-tailed *U* test. ^#^*p*<0.05.

Anti-inflammatory macrophages did not change in recruitment from days 1 to day 2; however, this subpopulation is 1.5-fold greater on endometriosis day 1 compared to control days 1 and 2 with no differences between endometriosis day 2 and days 1 and 2 in endometriosis with exogenous hormones (Figure 3D). The increase in anti-inflammatory macrophages in endometriosis day 1 maybe a compensatory response to regulate proinflammatory responses on day 1 in endometriosis. An additional subpopulation of anti-inflammatory macrophages was characterized by the presence of integrins (i.e., CD11b, CD11c), CD115, MHC class II (HLA-DR), and adhesive markers (i.e., CD54, CD31) with the absence of CD86 (Figure 3E). These cells did not differ from day 1 to day 2 in menstruation in control or endometriosis samples. The endometriosis samples from days 1 and 2 increased compared to control days 1 and 2 (3- and 2-fold, respectively) and not different from endometriosis with hormones days 1 and 2. This subpopulation also indicates a shift towards increased anti-inflammatory adhesive macrophages in endometriosis which may also be compensating to reduce a proinflammatory environment in women with endometriosis. Lastly, as many dendritic cells are monocyte derived, dendritic cells were examined and a trend of increased recruitment was seen from day 1 to day 2 in each condition (Figure 3F). These findings demonstrate the shift in increased dendritic cell recruitment occurs in all conditions; however, in the endometriosis with exogenous hormones group, day 2 increased by nearly 2.5-fold compared to control day 2, but was not different from endometriosis day 2. Overall, the differences in subpopulations between women with and without endometriosis suggests altered pro-inflammatory and anti-inflammatory subpopulations may contribute to shifts in the inflammatory environment in endometriosis.

### Adaptive immune subphenotyping reveals altered T cell population recruitment in endometriosis

Antigen presenting cells like macrophages and dendritic cells directly activate the adaptive immune system through interactions with T cells (47). Total T cells were <5% of the overall WBC population. Total T cells did not differ from day 1 to 2 in control women (Figure 4A). In women with endometriosis, T cells were approximately 2-fold higher on day 1 compared to control day 1, but not different from any other condition or day (Figure 4A). The total T cell data suggests an overabundance in endometriosis on day 1 of menstruation. Because T helper cells activate cytotoxic T cells and B cells (48), the presence of CD4 (T helper) and CD8 (T cytotoxic) were examined. In healthy controls, the level of T helper cells increases on day 2 of menstruation, but the levels in endometriosis on days 1 and 2 are greater by 50% compared to control day 1 (Figure 4B). Endometriosis day 1 and 2 T helper cells were not different than control day 2 or either day in the endometriosis with exogenous hormones group. These findings may suggest T helper cells are recruited earlier (i.e., in day 1) in women with endometriosis. Regulatory T cells, a subset of T helper cells, did not differ with condition or day across all groups (Figure 4C). Cytotoxic T cells did not differ by condition or day, but decreased 15% in day 1 of endometriosis compared to control day 1 and an ∼20% decrease in day 2 of endometriosis day 2 compared to control day 2 (Figure 4D). The decreased cytotoxic T cells in endometriosis day 1 and 2 could be due to decreased recruitment/activation by T helper cells suggesting a potential disruption in the activation mechanism of cytotoxic T cells.

**Figure 4:**
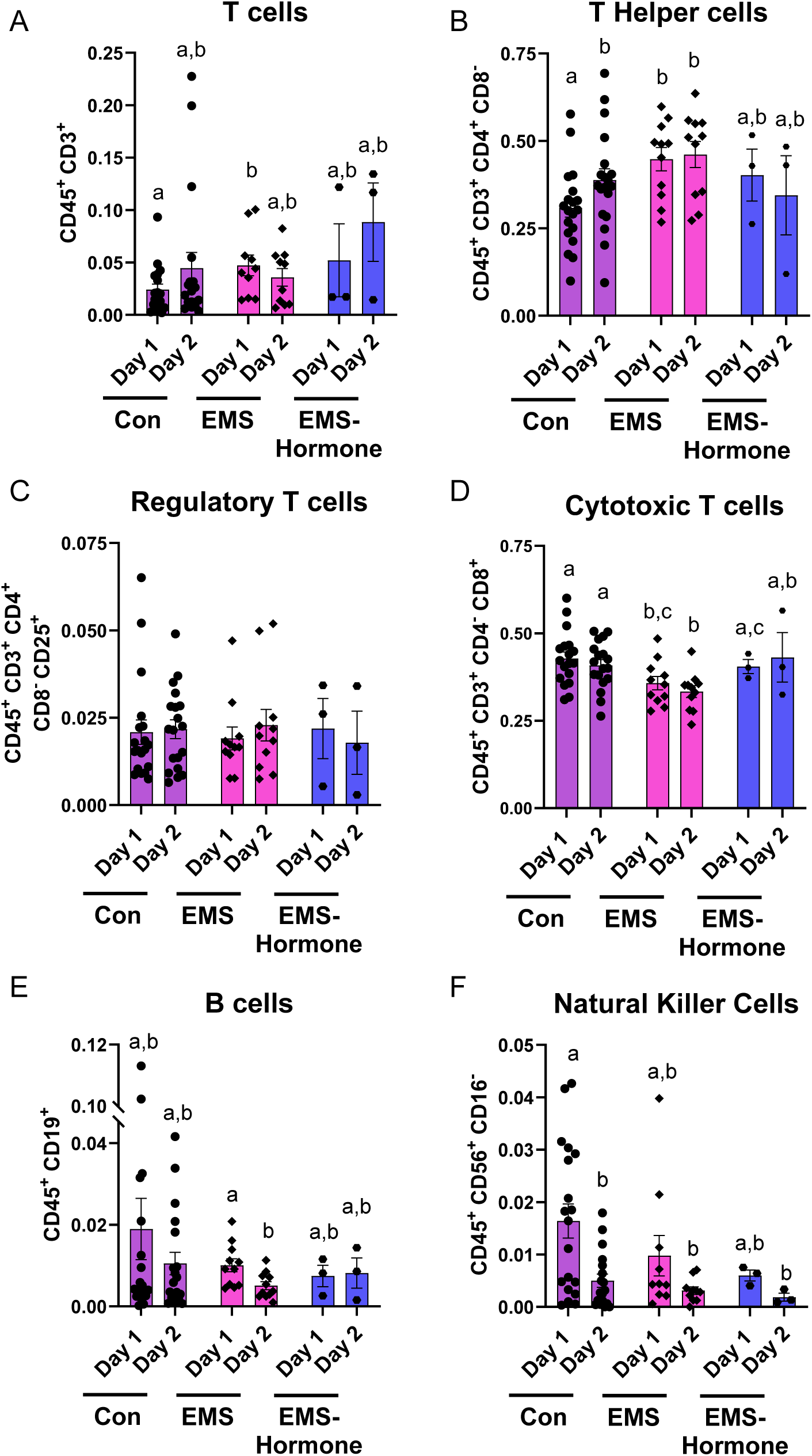
Women with endometriosis have increased T helper cells compared to control. **(A)** Quantitation of total T cells to total CD45^+^ ratio. Total T cells were defined by: CD45^+^ CD3^+^. **(B)** Subpopulation ratios were calculated to total T cells. T helper cells were defined as: CD45^+^ CD3^+^ CD4^+^ CD8^-^. **(C)** Regulatory T cells were defined as: CD45^+^ CD3^+^ CD4^+^ CD8^-^ CD25^+^. **(D)** Cytotoxic T cells were defined as: CD45^+^ CD3^+^ CD4^-^ CD8^+^. **(E)** Quantitation of total B cells to total CD45^+^ ratio. B cells were defined as: CD45^+^ CD19^+^. **(F)** Quantitation of total natural killer cells to total CD45^+^ ratio. Natural killer cells were defined as: as: CD45^+^ CD56^+^ CD16^-^. Control (*n*=19); endometriosis participants (*n* = 11); endometriosis with hormones (*n*=3). Data represent ± SEM. Statistical significance for each graph was determined by two-way ANOVA, followed by nonparametric, Mann-Whitney, 1-tailed *U* test. ^#^*p*<0.05.

We next examined B cells which are ∼2% of the overall WBC population in ME. The levels of B cells in healthy women did not differ from day 1 to day 2 (Figure 4E). In women with endometriosis, the levels of B cells did not differ from control women on day 1, but the levels decreased by ∼2-fold in endometriosis day 2 compared to day 1. Although the other conditions do not share this significant decrease between days 1 and 2, endometriosis may result in a decrease in the antibody release due to decreased B cell numbers on day 2. Uterine NK cells were examined in ME previously (1), but not by day of menstruation; therefore, NK (CD56^+^) cells were examined on days 1 and 2 of menstruation. A 3-fold decrease in control day 2 compared to control day 1 was observed in the NK cell populations (Figure 4F). However, the same decrease in recruitment was not observed between day 1 and 2 of the endometriosis or endometriosis with exogenous hormones indicating a dysfunction in the abundance of NK cells between days in endometriosis compared to controls. Altogether, the alterations in immune populations and recruitment differences found in endometriosis may contribute to immunological dysfunctions and endometriosis pathogenesis.

### Differentially expressed protein analysis of ME identified upregulated proteins for each day and condition

To further examine pathways of cellular action in ME, proteomics was done on the acellular portion of ME to identify differences based on condition and day. Samples were depleted of the top 14 abundant proteins and hemoglobin to capture less abundant proteins. Samples were analyzed with mass spectrometry for nested peptides and abundance levels. Due to sample size and variability observed in ME, a fold change of >1.5-fold was chosen. Pearsons correlation used to generate a heat map (Figure S3), revealed that control day 1 is most similar to control day 2, endometriosis day 1 is most similar to endometriosis day 2, control day 2 and endometriosis day 2 are somewhat similar, and finally, endometriosis day one is distinct from control day 1 and control day 2. Translationally, the paired days of each condition are similar to one another, but variances are found when comparing control day 1 to endometriosis day 1 and control day 2 and endometriosis day 2. Potential target proteins >1.5-fold different were examined by volcano plots. Over 1,000 total proteins were identified in ME, but not in all samples. They were filtered to ensure one of the four groups included that protein in >60% of the samples and the count dropped to 839. An additional 267 proteins were eliminated because they did not differ by condition or day. Finally, 572 proteins were used to generate volcano plots based on comparisons between the paired days (day 1 vs day 2) of each condition and the same day but different conditions (control day 1 vs endometriosis day 1) (Figure 5A-D).Fewer proteins overall were identified that increased in endometriosis samples compared to control samples (control day 1 vs endometriosis day 1 with 197 vs 143, control day 2 vs endometriosis day 2 with 195 vs 107) suggesting that the control samples are more active than endometriosis samples.

**Figure 5.**
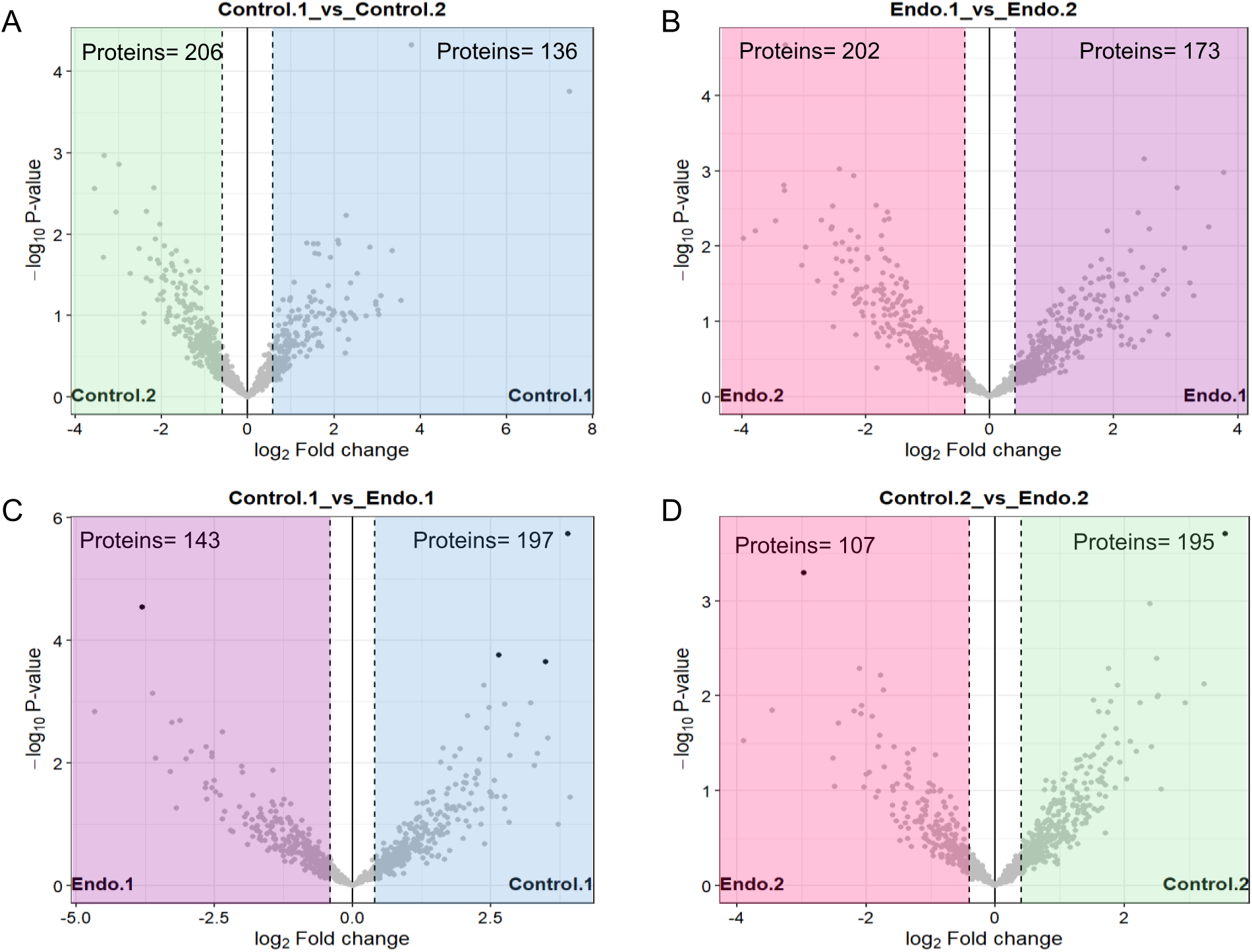
Differentially expressed proteins are more abundant in control ME. Volcano plots were generated from differential expression protein analysis with biological relevance determined by >1.5-fold. The number of proteins identified in each condition and day for each comparison is located in the upper corners. **(A)** Control day 1 (blue) compared to control day 2 (green). **(B)** Endometriosis day 1 (purple) compared to endometriosis day 2 (pink). **(C)** Control day 1 compared to endometriosis day 1. **(D)** Control day 2 compared to endometriosis day 2. Control day 1 and 2 (n=6), Endometriosis day 1 (n=5), Endometriosis day 2 (n=6).

### Differentially expressed protein analysis of ME demonstrates alternative activation and regulation of cellular pathways

Based on these comparisons, the identified proteins were further characterized to reveal pathways of interest using the STRING database (49). The protein lists generated from each comparison above was entered into the STRING database for functional enrichment analysis. The pathways were selected based on relevance to the process of menstruation and on the protein count or number of proteins associated with individual pathways. The pathways were then grouped by strength and similarity of the proteins associated with the selected pathways. The first comparison was between control days 1 and 2 to establish a baseline of which pathways were activated in each day (Figure 6A and B). Control day 1 compared to control day 2 showed similar pathways, but control day 2 lacked catabolic activity pathways and suggests that catabolism is prioritized on day 1, but not day 2 of menstruation in controls. The next comparison was to establish the pathways that were occurring in endometriosis day 1 and 2 (Figure 6C and D). Similar pathways were identified in each day in endometriosis, but the strength of cell adhesion molecule binding pathway was stronger in endometriosis day 2 indicating this pathway is more active in day 2. Interestingly control day 1 and endometriosis day 1 (Figure 6A and C) show the same pathways in a similar organization; however, endometriosis day 1 has less cell adhesion binding molecules than control samples. Control day 2 and endometriosis day 2 show similar pathways when compared with their day 1 of the same condition (Figure 6B and D). However, the hydrolase pathway for metabolism and cellular signaling were not present in the endometriosis day 2 samples when compared to endometriosis day 1 samples (Figure 6D). Once the base functions of each condition with their respective paired day was established, control day 1 compared to endometriosis (Figure 6E and F) pathways were similar to the previous results for day 1 in both conditions. When control day 1 was compared to control day 2 and endometriosis day 1 was compared to endometriosis day 2, similar pathways were identified in both control day 1 and endometriosis day 1 as well as control day 2 and endometriosis day 2. When control day 1 and endometriosis day 1 were compared, different proteins were identified to be upregulated for each condition; however, the same pathways were still identified indicating that both control and endometriosis are conducting similar processes, but based on the enrichment, the upregulation of different proteins and generation of the same pathways suggest an alternative activation or processes related to these pathways. Interestingly, control day 1 and endometriosis day 1 did not show any protein differences in hydrolase function indicating that this function is likely similar in day 1 of both conditions. The last comparison performed was control day 2 compared to endometriosis day 2 (Figure 6G and H). Endometriosis had similar pathways to control day 2; however, the number of proteins associated with each immune pathway (e.g., neutrophil degranulation, secretory granule, and innate immune system) contained lower numbers of associated proteins in the pathway which indicates that endometriosis day 2 is communicating with the immune system, but in a different manner than what occurs in control day 2. The change in protein numbers further suggests altered immune function in endometriosis via differentially regulated pathways or the use of alternative pathways compared to control samples. Additionally, endometriosis day 2 compared to control day 2 did not show pathways for peptidase activity, catabolic processes, and hydrolase activity indicating that these pathways are more active or increased in control day 2 compared to endometriosis day 2 proposing that endometriosis day 2, having the lowest number of proteins identified, is less active or not activating the pathways needed. Endometriosis day 2 compared to control day 2 (Figure 6G and H) did not have peptidase activity and catabolic processes, in contrast, endometriosis day 2 does have these associated pathways when compared to endometriosis day 1 (Figure 6D); thus, endometriosis day 2 is less active in these processes than control day 2, the opposite of control. Overall, the proteomics confirmed pathways are altered and functional differences exist between endometriosis and control samples that likely lead to altered inflammatory processes in the pathogenesis of endometriosis.

**Figure 6.**
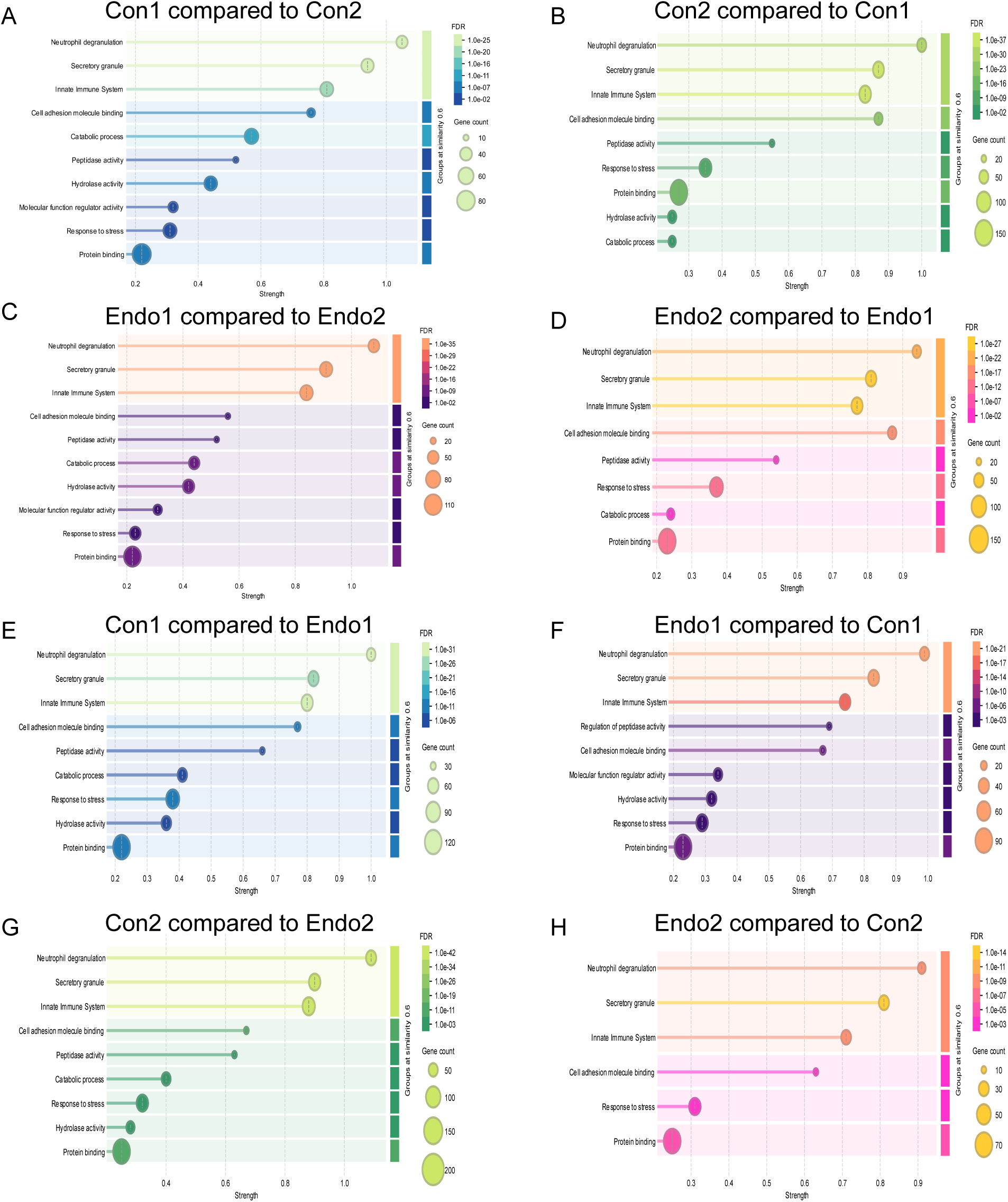
Selected enrichment pathways from STRING database demonstrates altered functional pathways between control and endometriosis. Proteins generated based on 1.5-fold change were analyzed using STRING database to elucidate potential functional pathways comparing condition and day. Protein pathways are grouped based on similarity >0.6 and organized by strength of pathway connection. The dot size indicates the number of proteins associated with a particular pathway and color indicates the false discovery rate. **(A)** Control day 1 (blue color scale) associated protein pathways compared to control day 2. **(B)** Control day 2 (green color scale) associated protein pathways in comparison to control day 1. **(C)** Endometriosis day 1 (purple color scale) associated protein pathways compared to endometriosis day 2. **(D)** Endometriosis day 2 (pink color scale) associated proteins compared to endometriosis day 1. **(E)** Control day 1 associated protein pathways compared to control day 2. **(F)** Endometriosis associated protein pathways compared to control day 1. **(G)** Control day 2 associated protein pathways compared to endometriosis day 2. **(H)** Endometriosis day 2 associated pathways compared to control day 2. Con=control, Endo=endometriosis. Control day 1 and 2 (n=6), Endometriosis day 1 (n=5), Endometriosis day 2 (n=6).

## Discussion

The functions and mechanisms of the immune system in endometriosis lesion formation and pathogenesis are not well understood (13, 43, 50, 51). We and others have shown neutrophils and associated factors are an effector cell in the initiation and pathogenesis of endometriosis (43, 52–54) and likely direct the recruitment and function of other WBCs (55–59). Although endometriosis is a complex disease with multiple factors influencing the progression of disease, the immune system is proposed as a leading contributor to tolerance and attachment of endometrial tissue in the peritoneal cavity as a result of retrograde menstruation. Utilizing ME to evaluate and characterize WBC populations is a non-invasive method to sample the cell types which reflux into the peritoneal cavity during menstruation. The influx of WBCs and endometrial tissue into the peritoneal cavity has the potential to establish new lesions and exacerbate the local inflammatory environment every month. Since only 10% of women develop endometriosis and they have increased associations with autoimmune diseases (29, 30), the immune system is a driving factor in disease pathogenesis. Our study, to our knowledge, is the first to deep immunophenotype the WBCs found in ME. We provide evidence women with endometriosis have altered innate and adaptive immune subpopulations and variability of immune subpopulations increases with exogenous hormonal therapies. In contrast, shifts that occur in cell population numbers in control day 1 and 2 samples are not always present in endometriosis participants, implicating immune dysfunction. Furthermore, our study identifies potential alternatively activated functional pathways and differentially regulated protein pathways in ME of women with endometriosis supporting the hypothesis that the immune system and inflammatory environment in women with endometriosis is functionally different than what occurs in normal menstruation.

Our findings regarding symptomatology and infertility reproduce differences already established between healthy control and endometriosis groups. Although not directly associated with the immune system, the distinct increase in pain related symptoms in endometriosis participants indirectly suggests menstruation is more inflammatory leading to the increase in symptom severity. Immune cells like neutrophils and macrophages are implicated as effector cells contributing to pain responses chronic diseases (15, 18, 60, 61); thus, they are likely contributing to amplifying pain symptoms in endometriosis. Additionally, the increase in symptom severity in women with endometriosis may be linked to the increased menstrual flow rate suggesting an additional contributing factor that may increase retrograde menstruation in endometriosis.

The striking discovery of a morphologically distinct phenotype of hypersegmented nuclei and the abundance of vacuoles in neutrophils in ME from women with endometriosis uncovers an avenue for endometriosis to be diagnosed noninvasively using a straightforward isolation, stain, and visualization that can be done economically. The altered cell morphology denotes an aged phenotype and physiologically represents differed physiological function(s) of the cell, their responses to inflammation, and/or transcriptional changes resulting in the modified nuclear arrangement. To date, this particular phenomenon has not been elucidated in the ME of women with endometriosis; thus, these novel findings position ME as a tool to diagnose endometriosis. Our subject population was homogenous and primarily Caucasian; however, if a ME test can be done economically, we foresee women from socioeconomically diverse groups to now acquire a diagnosis of endometriosis.

Since neutrophils are recruited prior to the onset of menstruation to prime the uterine microenvironment for the inflammatory event of menstruation (23), our findings suggests a dysfunction in the uterine microenvironment or in the neutrophils themselves from women with endometriosis that shift the neutrophils toward an aged phenotype. A strength to the study design is the inclusion of days 1 and 2 and clarification on the exact day of menstruation to evaluate the immune cell abundance. Having samples from day 1 and day 2 increases the depth of knowledge of what occurs normally in menstruation. We uncovered that neutrophils are the most abundant cell type present in ME as is found in peripheral blood. Having both days 1 and 2 of menstruation allowed us to uncover shifts in cell abundance/recruitment and, for example, allowed us to uncover that aged neutrophils are typically increased on day 1 of menstruation compared to healthy women and endometriosis day 2. Although the lifespan of neutrophils is short in comparison to other cells (62, 63), they are instrumental in clearance of endometrial tissue and recruitment of phagocytic cells to further remove debris for the resolution and repair of the endometrial lining (23, 64). Whether the influx of neutrophils prior to menstruation causes an alternative priming of neutrophils to prematurely age in endometriosis or whether they are recruited to the uterus in an already aged state is unknown; however, the findings support a dynamic difference in the neutrophils found in women with endometriosis.

After neutrophils, macrophages are the next abundant cell type in ME and they are directly and indirectly influenced by neutrophils as a result of cytokine release (65). We find antigen presenting proinflammatory macrophages less in endometriosis participants than healthy participants on day 1 of menstruation, but they are recruited at the same levels in day 2. In contrast, endometriosis day 1 has more anti-inflammatory macrophages than control day 1 indicating endometriosis ME is compensating with an anti-inflammatory response instead of a proinflammatory response likely in response to the increased inflammation generated from the neutrophil subpopulations; thus, furthering the immune dysfunction as the anti-inflammatory macrophages are not able to mitigate the inflammation. The predominant presence of foamy macrophages in ME from women with endometriosis further supports the hypothesis that anti-inflammatory macrophages are functioning to mitigate the overly inflammatory microenvironment. Additionally, an adhesive anti-inflammatory macrophage was more abundant in endometriosis ME on both days compared to control, again supporting alternative compensation or mitigation of the proinflammatory responses in patients with endometriosis. Furthermore, a decreased pro-inflammatory phenotype along with an increased anti-inflammatory phenotype indicates an imbalance in the subpopulations of macrophages in women with endometriosis is altering the process leading to decreased clearance of ME to form endometriotic lesions and/or shifting the microenvironment to alter the inflammatory response that contributes to pain symptomatology.

As antigen presenting cells, macrophages and dendritic cells stimulate T cells to further resolve injury or an inflammatory event (66, 67). Increased total T cells in endometriosis day 1 compared to control day 1 indicates T cells are recruited to the endometrium earlier in endometriosis ME than in control ME. Interestingly, the T helper cell numbers on days 1 and 2 in endometriosis are similar to the control day 2 but not control day 1 indicating the T helper response in endometriosis more closely resembles the control day 2 phenotype. Interestingly, a specific subset of helper T cells (T_h_17 cells) are elevated in peripheral blood and in the endometrial tissue of endometriosis patients (20) corroborating the increase in T helper cells found in endometriosis ME. Of note, the number of cytotoxic T cells was less in endometriosis ME compared to control ME, again, supporting previous reports of reduced cytotoxic T cells and their function in endometriosis (16, 57). Lastly, B cells are fewer in endometriosis day 2 indicating they are either not recruited and/or activated in day 2 like in day 1 because of the disconnect in the resolution responses and cellular communication seen in ME from women with endometriosis. Collectively, in endometriosis the altered recruitment of various WBC subpopulations indicates signaling or recruitment dysfunctions in endometriosis, ultimately disrupting the synchrony of the inflammatory process to exacerbate and/or dampen the normal inflammatory responses of menstruation.

The acellular fraction of ME provides evidence of additional physiological dysfunctions in the ME of women with endometriosis. During menstruation, inflammatory processes induce apoptosis and the release of neutrophil extracellular traps (NETs) which releases cfDNA (46, 68). Although phagocytic cells clear cfDNA and cellular debris (45, 46), cell free DNA accumulation may be a transient indicator of clearance, NETs, or apoptotic activity. Even with variability in ME from individual physiological diversity, cell free DNA release increased from day 1 to day 2 in control ME and was not observed in endometriosis ME, indicating a dysfunctional clearance of cellular debris and/or less cellular processes to release cfDNA. Although the sample size is low and ME has inter- and intra-variability, proteomic analysis provided prospective pathways of target for future studies. The enrichment analyses identified variances in pathway function between control and endometriosis for condition and day of menstruation. In general, the proteins in ME were highly specific for immune pathways which supports its important role during menstruation. Of note, similar pathways were identified in control and endometriosis, but different proteins from the same overall pathways were identified to be increased in each condition suggesting alternative activation or mechanistic dysregulation of the same pathways in endometriosis compared to control. The altered signaling is largely associated with metabolism and cell signaling, crucial responses needed for immune cell activation (69). The strength in separating days 1 and 2 uniquely provided insights into the normal shifts in cell activity and recruitment which would be missed during random collections or from combining multiple days of ME.

Collectively, our study provides further evidence of immunological dysfunction in endometriosis, but more specifically how the dysfunction during menstruation may contribute to the pathobiology of endometriosis. By defining the normal range of immune cells and protein abundances, we were able to characterize immunological shifts in subpopulations of neutrophils, macrophages, and lymphocytes, as well as alternatively activated protein pathways occurring in the ME of women with endometriosis. While a limitation to our study is the smaller sample size and lack of ethnic diversity, the analysis provides evidence of biologically relevant immune dysfunctions for deeper understanding of endometriosis pathology. Presently, non-white ethnicities encounter increased risk of perioperative and postoperative complications with laparoscopy (70, 71) in addition to financial burdens from lack of insurance coverage (72); thus, our findings leading to a noninvasive diagnostic could reduce socioeconomic and health disparities by improving equity and improved access to an endometriosis diagnoses for previously underrepresented populations. Furthermore, as a result of the difficulty to invasively acquire endometriosis lesions, peritoneal fluid, uterine biopsies, and peripheral blood in both endometriosis and matched control samples, our study supports ME can provide a non-invasive method to study endometriosis pathogenesis. Ultimately, our findings uncover that ME can be used as a tool to diagnose endometriosis—decreasing the need for surgery, the 7-9 years for a diagnosis, the economic burden, and the socioeconomic disproportion in having a diagnosis.

## Methods

### Study approval, ME collection, and collection

The study protocol was approved by the Institutional Review Board (IRB) at University of Cincinnati (IRB No. 2015-7749). Only females were part of our study design, because endometriosis is a gynecological disease that occurs in people who menstruate (i.e., genetically XX). Patients gave written informed consent prior to participation in the study. ME was collected for 6-24 hours on days 1 and 2 of menstruation using a DIVA™ cup. Day 1 was characterized by active bleeding, not spotting. Once collected, samples were refrigerated and transported on wet ice. Eligible participants were naturally menstruating, not pregnant, aged 21–44, did not have an intrauterine device, had no use of hormone contraceptives for at least 3 months prior to collection for the endometriosis and healthy groups, had no history of cancer, had no active infections, and were not currently on antibiotics. Patients with endometriosis were laparoscopically confirmed. Patient demographics were as follows: Healthy (3 African American/Black, 3 Asian, 13 White; age 25–44 years), Endometriosis (11 White, age 21–40 years), endometriosis with Hormones (3 White; age 27-40). ME was filtered through sterile gauze and centrifuged for 30 minutes at 277*g* to separate acellular and cellular fractions. The acellular fraction was saved at −80◦C for proteomics analysis. Cells were resuspended in PBS + 0.5% BSA + 2 mM EDTA buffer (FACS buffer), spun, and filtered through 70 μm and 40 μm filters to remove epithelial/stromal cells. Immune cells were isolated via density centrifugation composed of equal parts Histopaque (11191, Sigma-Aldrich) and LymphoprepTM (07851/07861, STEMCELL Technologies) for 20 minutes at 277*g*. The leukocytes were removed, washed, and resuspended in FACS buffer, then utilized for immunophenotyping using spectral flow cytometry and differential staining. Due to variability in menstrual flow and hours collected, samples were normalized per mL of fluid and the ratio was determined by the number of cell population (i.e. neutrophil) in each subpopulation per mL corresponding to the total number of CD45^+^ cells per mL.

### Flow Cytometry Analysis

The WBCs were isolated and resuspended in FACS buffer. In each well, 2 million cells were plated, blocked with 0.5% normal rat serum + 0.5% normal mouse serum + 0.004% anti– human Fc receptor binding (14-9161-73, Invitrogen) in FACS buffer for 30 minutes, and then stained for cell surface markers identifying neutrophils. CXCR and CCR markers were first added, one at a time for 10 minutes each, before adding the additional cell surface markers as a mixture for 30 minutes (Supplemental Table 1-2). In the antibody mixture, 10 µL of Super Bright Complete Staining Buffer (#SB-4401-75, Invitrogen) and 25 µL of CellBlox™ Blocking Buffer was included. After antibody staining, cells were fixed with 1% paraformaldehyde for 5 minutes. Stained cells were analyzed using spectral flow cytometer on the FACS Aurora (BD Biosciences). Single-stained controls for each cell surface marker were used to validate antibodies, spectral unmixing, and compensation using FlowJo software (Verizon 10.10) for analyses.

### Deep immunophenotyping of ME gating strategies

As deep immunophenotyping, to our knowledge, has not been done on the WBCs from ME, to characterize these morphological differences observed in the differentials, the WBCs isolated from ME were deep immunophenotypes via spectral flow cytometric analysis with advanced gating strategies (Figure S2). Gating began with a “Time gate” to eliminate and/or detect flow rate disturbances, then a gate was formed around the location of the population of interest (i.e., granulocyte, macrophage, lymphocyte). Since WBC subpopulations share many surface markers due to shared functions or activation status, gates were placed in the expected locations for the population of interest based on forward and side scatter heights (73, 74) to reduce the potential for false discovery. Single cell gates (i.e., SSC-A and SSC-H) and CD45^+^ (i.e., SSC-H and CD45^+^) gates were drawn prior to gating on cell specific markers. Granulocytes were gated with a pan-marker of CD66b with CD193 as the exclusion marker used for eosinophil activation (i.e., CD66b^+^ CD193^-^) (Figure S2A). Based on differential images, eosinophils are minimal in abundance; thus, this marker was selected for eosinophil exclusion. Neutrophils, when mature, express CXCR2 (75) and, when aged, express CXCR4 (38, 76); therefore, they were gated on CD16 and then CXCR2 and CXCR4 for subsequent gating for maturity and activation of subpopulations.

Macrophages were also gated based on forward and side scatter heights (i.e., FSC-H and SSC-H) (73, 74) to reduce the potential for false discovery of other populations of cells. From the known locations, single cell gates (i.e., SSC-A and SSC-H), and CD45^+^ (i.e., SSC-H and CD45^+^) gates were drawn prior to gating on cell specific markers. They were initially gated for CD14 and CD16. CD16 positive (CD16^+^) indicates a pro-inflammatory phenotype while CD16 negative (CD16^-^) is associated with an anti-inflammatory phenotype (Figure S2B). Subphenotypes of macrophages were determined by subsequent gating from these initial gating strategies.

The smaller populations of WBCs present in ME were gated in a similar manner as neutrophils and macrophages. For dendritic cells, a CD14 low (CD14^lo^) population was utilized for gating. T cells were gated initially on CD3 and subsequently on population specific markers (i.e., CD4^+^, CD8^+^, etc). B cells were identified as CD19^+^ and natural killer cells were identified as CD56^+^ (Figure S2C). These gating strategies were utilized to reduce potential false positive populations and identify specific surface marker differences between healthy controls and endometriosis groups.

For all samples in the analyses, to normalize the data for collection time and sample volume variability, WBC populations were first calculated per mL of ME collected. Next, a ratio was calculated for the main cell types (i.e., neutrophils, macrophages, T cells, B cells, NK cells, dendritic cells, and eosinophils) by dividing the total parent cell population (i.e., neutrophil alone, macrophage alone) per mL by the total CD45^+^ cells per mL to understand the composition of each immune cell type in ME in each sample collected. Subphenotype ratios were calculated by dividing the subpopulation calculation per mL by the primary cell population compared to total CD45^+^ cells to understand the abundance of subpopulations in ME.

### Cytospin and Differential Staining

Immune cells isolated from ME were used for differential staining. Cells were counted (100k and 75k) and cytospun onto slides in 100 μL of in PBS + 0.5% BSA with a Thermo Shandon Cytospin 4 machine for 5 minutes at 400 rpm. Slides were stained with a modified Giemsa stain according to manufacturer’s instructions.

### Cell free DNA Isolation

A 250 µL aliquot of the acellular fraction of ME was utilized for cell free DNA isolation. Samples were diluted to a total volume of 3 mL with nuclease-free water due to the thick viscosity of the ME. The cfDNA was isolated using a Quick-cfDNA/cfRNA Serum and Plasma kit (#R1072, Zymo Research) following the manufacturer’s protocol.

### Proteomics Analysis

Aliquots of the acellular fraction of ME were utilized for proteomics. Hemoglobind (H0145-05, Biotech Support Group) was used per manufacturer’s protocol to deplete hemoglobin. The top 14 most abundant proteins in blood products were removed using High Select Depletion Spin columns (A36369, ThermoFisher) following manufacturer’s protocol. Samples were analyzed further by the University of Cincinnati’s Proteomics Core facility using tandem mass spectra in combination with high confidence peptide identifications to generate protein with nested peptide abundance levels. The abundance data was further analyzed using Rstudio (version 2024.12.1+563) with installed packages: Biocmanager, DEP, dplyr, ggplot2, limma, tidyr, tibble, ggfortify, and enhanced volcano. Differential protein analysis and enrichment was generated using the DEP package to generate log2fold values for comparisons between each group and condition (Disease, Day). A 1.5-fold change cutoff was used. For enrichment analysis, the STRING database (49) was used to generate potential pathway interactions based on protein lists generated through differential protein expression analysis. The enrichment criteria were based on high confidence and pathways were selected by gene count and strength.

### Statistics

When comparing 3 or more groups for multiple comparisons, nonparametric data were analyzed using a Kruskal-Wallis test with the mean rank of each column compared with the mean rank of every other column. If the Kruskal-Wallis test was significant (*p*<0.05), then Mann-Whitney, 1-tailed U tests were performed for validation. Statistical analyses were performed using GraphPad Prism version 10.2.0 (GraphPad Software). When comparing 2 groups, nonparametric, Mann-Whitney, 1-tailed U tests were performed. For multiple-group comparisons, means not sharing a letter are significantly different from each other (*p*<0.05) and means sharing the same single letter or a letter in combination with other letters are not significantly different from each other. For U tests, the number symbol (#) indicates significant differences (*p*<0.05).

## Acknowledgements

All flow cytometric data were acquired using equipment maintained by the Research Flow Cytometry Facility in the Division of Rheumatology at CCHMC supported by NIH S10OD025045. We thank the assistance of the Research Flow Cytometry Facility and the guidance of Ken Quayle. We are appreciative of DIVA Cup for donating the DIVA Cups and to each participant for providing menstrual fluid. Mass spectrometry data were collected and analyzed in the UC Proteomics Laboratory under the direction of KD Greis, PhD. Funding for the Thermo Orbitrap Eclipse nanoLC-MS/MS system was obtained in part through an NIH high end instrumentation grant (S10OD026717-01). Sources of support to KAB include R01 HD097597 and University of Cincinnati College of Medicine Startup.

## Author Contributions

AS, SYT, KR, and KAB designed the immunophenotyping panels. PLD isolated and purified acellular proteins for proteomics. TRW, SAM, and KAB collected and isolated ME WBCs for spectral flow cytometry and differential staining. CFM helped optimize the collection of ME. SV and TRW wrote the code for proteomic analysis. TRW wrote the first draft of the manuscript and was edited by KAB. Analysis of data was performed by TWR, KAB, and PLD. KAB designed and developed the study. Each author read and made corrections to the manuscript.

## Supplemental Material

**Figure Supplemental 1.**
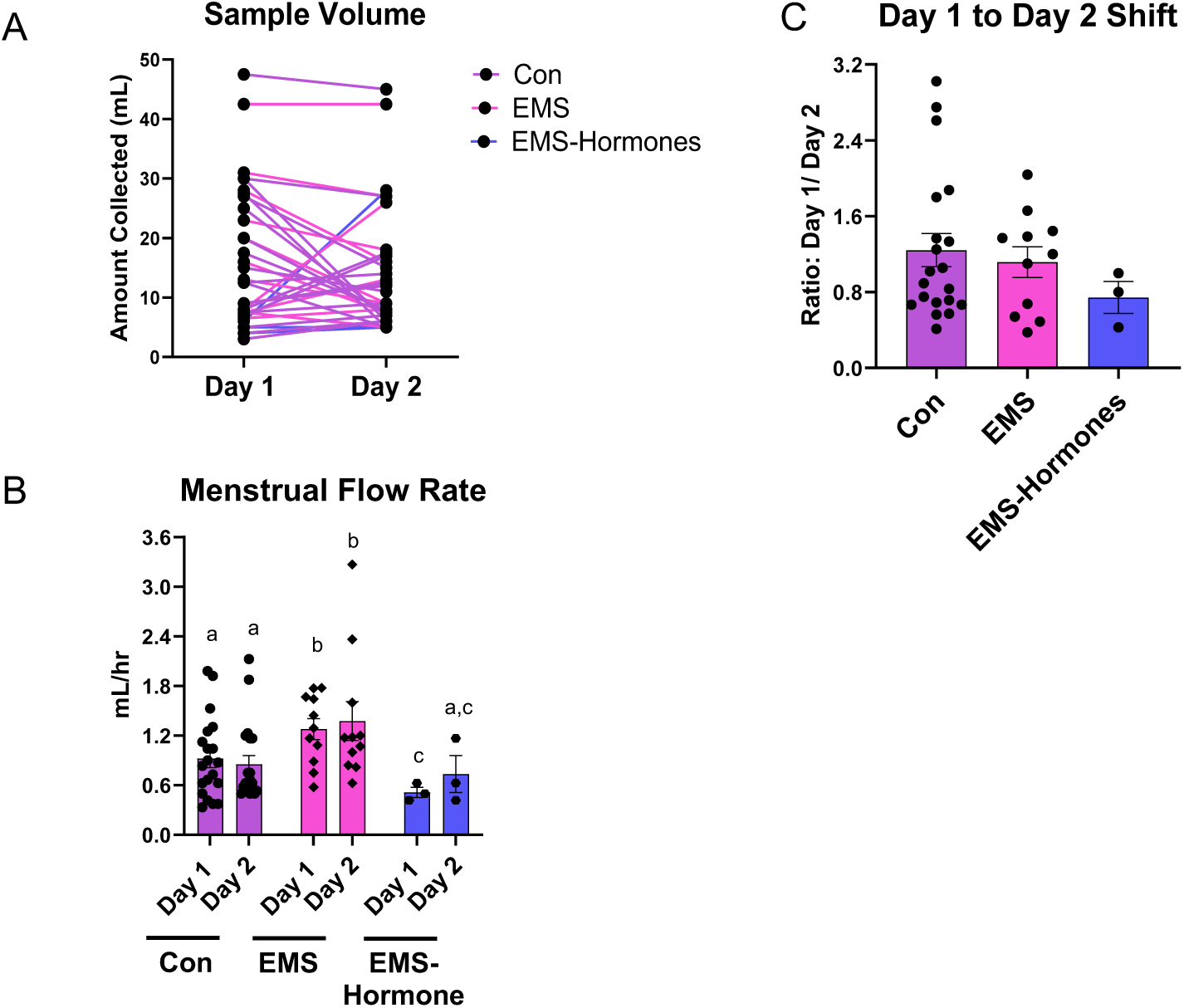
The menstrual effluent flow rate is increased in endometriosis day 1 and 2 samples. **(A)** Day 1 and 2 sample volumes collected from participants. **(B)** Menstrual effluent flow rate for each condition and day. **(C)** Menstrual effluent sample volume per hour ratio (day1/day2). Control (n=19), Endometriosis (n=11), Endometriosis with Hormones (n=3). Data represent ± SEM. Statistical significance for each graph was determined by nonparametric, Kruskal-Wallis test followed by Mann-Whitney, 1-tailed U test. p < 0.05.

**Supplemental Figure 2.**
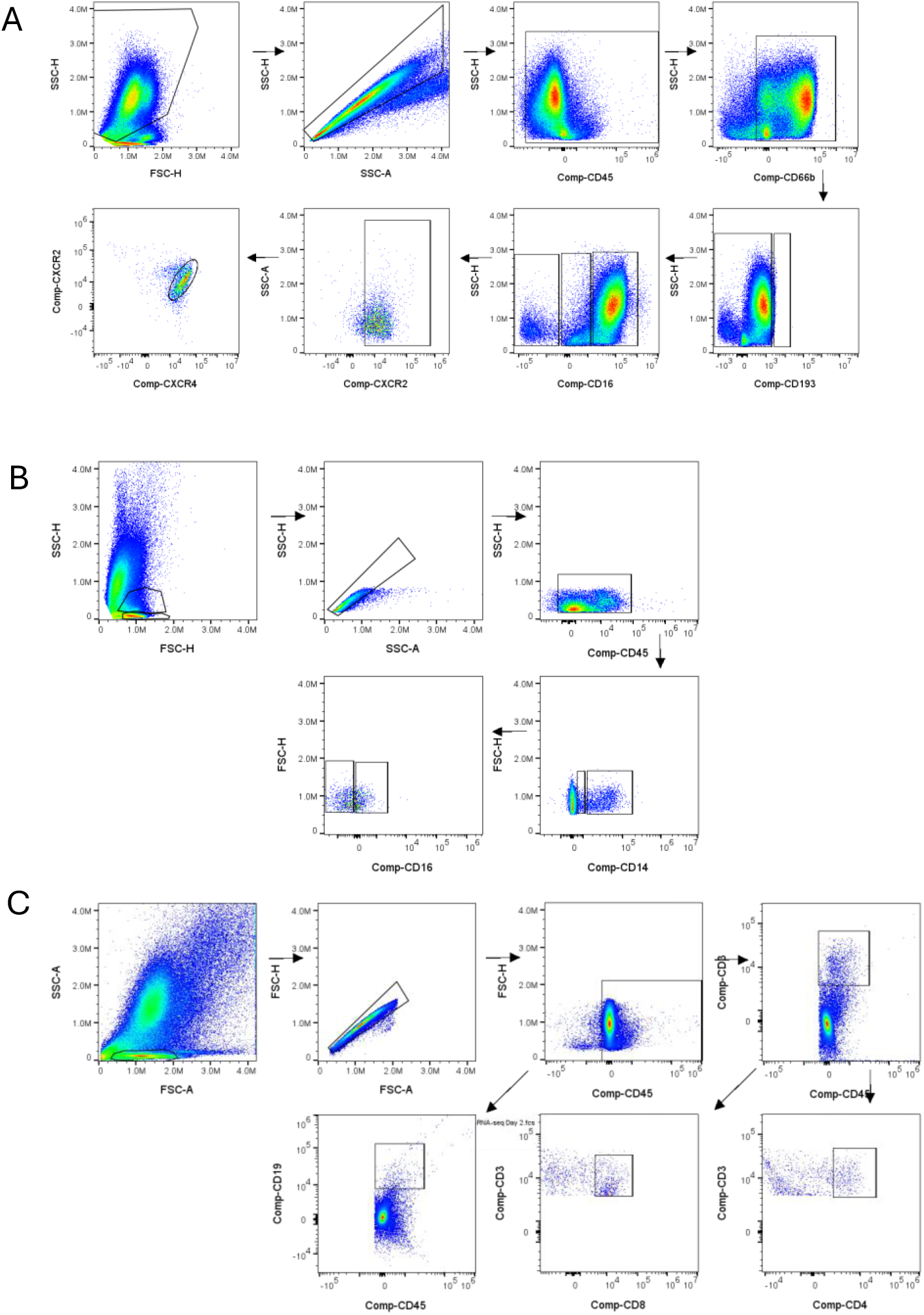
- Gating strategy of total WBCs isolated from human menstrual effluent. Cells were gated on time, population of interest, single cells. **(A)** Neutrophil initial gating strategy begins with CD45^+^, CD66b^+^, and CD193^−^ to identify total neutrophils with CD193^+^ used to identify eosinophils. **(B)** Macrophage grating strategies begin with CD45^+^, CD14^+^, CD16^+^ with the CD14^lo^ gate used to identify dendritic cells. **(C)** Lymphocyte gating strategy begins with CD45^+^, CD3^+^ and B cell CD45^+^, CD19^+^.

**Supplemental Figure 3.**
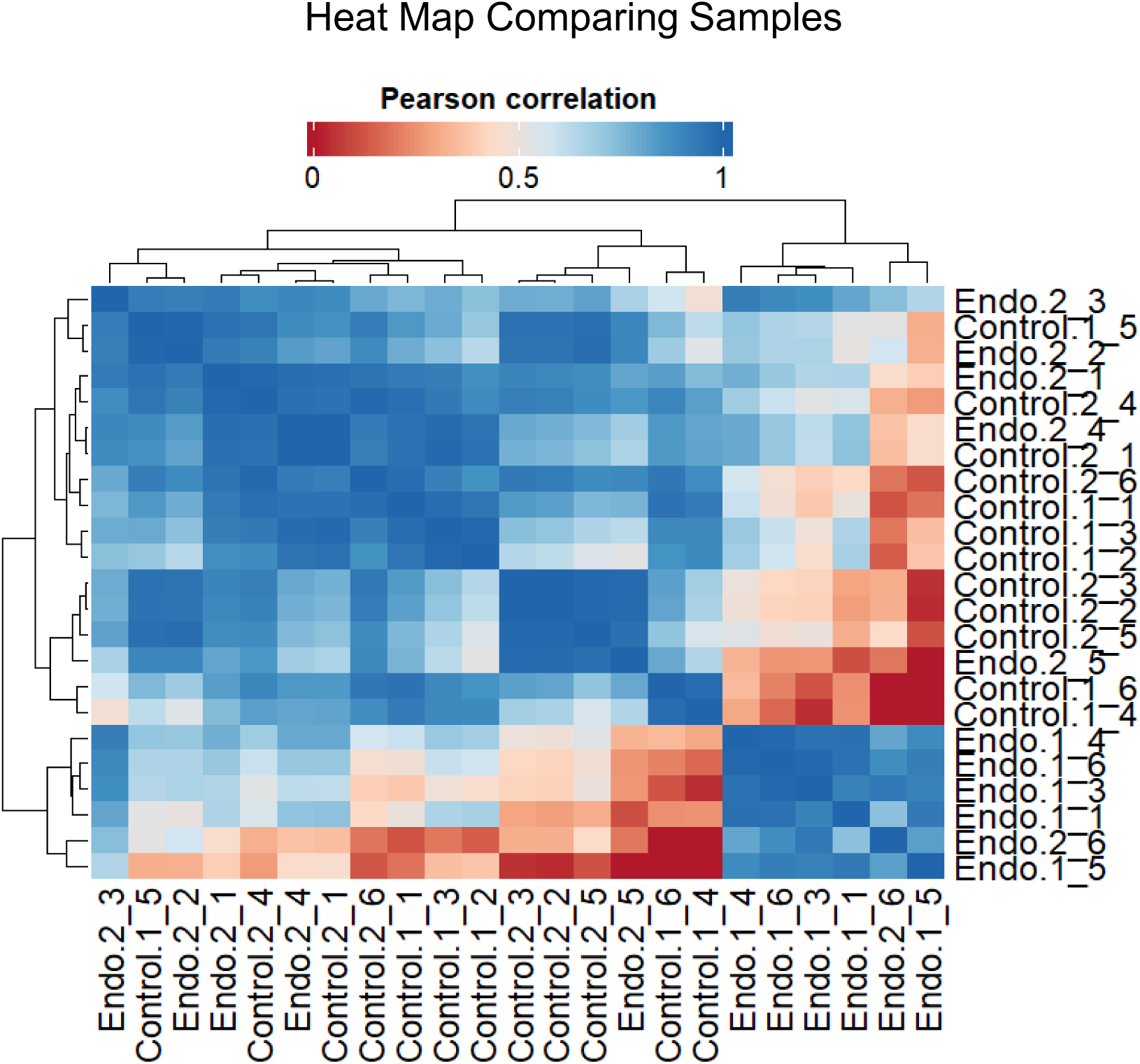
- Endometriosis day 1 proteomic samples are less similar to control day 1 and day 2 samples. Heat map of condition and day clustered by similarity using Pearson correlation. Blue indicating more similarity and red indicating less similarity between samples. Sample names are interpreted as condition.day-sample number. Control day 1 and 2(n=6), Endometriosis day 1 (n=5), Endometriosis day 2 (n=6).

**Supplemental Table 1.** Panel A. 27-color spectral flow cytometry panel for identification of neutrophil subtypes, eosinophils, lymphocytes, and endometrial stromal stem cells.

| Panel A |  |  |  |  |  |  |
| --- | --- | --- | --- | --- | --- | --- |
| Marker | Dilution (μl) | Fluorophore | Laser | Company | Catalog Number | Clone |
| CD25 | 1.25 | BUV563 | UV9 | BD | 612919 | 2A3 |
| CD64 | 1.25 | BUV737 | UV14 | BD | 612777 | 10.1 |
| CD196 (CCR6) | 2.5 | BUV496 | UV7 | BD | 612948 | 11A9 |
| CD11b | 0.31 | eFluor506 | V7 | Thermo | 69-0112-80 | M1/70 |
| CD90 | 2.5 | SuperBright436 | V2 | Thermo | 62-0909-42 | eBio5E10 |
| CD49d | 2.5 | BV480 | V5 | BD | 566134 | 9F10 |
| CD34 | 2.5 | SuperBright 645 | V11 | Thermo | 64-0349-42 | 4H11 |
| CD18 | 2.5 | SuperBright 600 | V10 | Thermo | 63-0189-41 | 6.7 |
| CXCR3 | 2.5 | BV785 | V15 | Biolegend | 353737 | G025H7 |
| CXCR1 | 1.25 | BV421 | V1 | BD | 743419 | 5A12 |
| CD62L | 0.5 | Pacific Blue | V3 | Thermo | MHCD62L28 | DREG-56 |
| CD45 | 0.5 | NFB610 | B6 | Thermo | H005T02B05 | 2D1 |
| CD3 | 1 | NFB660 | B7 | Thermo | H002T02B07 | UCHT1 |
| CD193 (CCR3) | 2.5 | AF488 | B2 | Thermo | 53-1939-41 | 5E8-G9-B4 |
| CD4 | 2.5 | NFB555 | B3 | Thermo | H001T02B03 | SK3 |
| CXCR2 | 5 | PerCP-eFluor710 | B10 | Thermo | 46-1829-42 | 5E8-C7-F10 |
| CD274 (PDL1) | 2.5 | PerCP-Cy5.5 | B9 | Biolegend | 329738 | 29E.2A3 |
| CD66b | 1.25 | PE-Cy7 | YG9 | Thermo | 25-0666-41 | G10F5 |
| CD19 | 2.5 | NFY660 | YG4 | Thermo | H004T03Y04 | 1D3 |
| CXCR4 | 2.5 | PE-eFluor610 | YG3 | Thermo | 61-9999-42 | 12G5 |
| CD8 | 1 | NFY690 | YG6 | Thermo | H003T02Y05 | OKT8 |
| CD11c | 1.25 | PE-Cy5.5 | YG7 | Thermo | 35-0116-42 | 3.9 |
| IGF1R | 5 | PE | YG1 | Thermo | MA5-16465 | 1H7 |
| CD16 | 0.5 | NFR700 | R3 | Thermo | H006T02B03 | 3G8 |
| CD31 | 0.63 | APC-eFluor780 | R7 | Thermo | 47-0319-41 | WM59 |
| CD54 (ICAM-1) | 5 | AF700 | R4 | Thermo | 56-0549-42 | HA58 |
| VEGFR1 | 10 | APC | R1 | RnD | 49560 | FAB321A |

**Supplemental Table 2.** Panel B. 27-color spectral flow cytometry panel for identification of macrophage subtypes, dendritic cells, and uterine natural killer cells.

| Panel B |  |  |  |  |  |  |
| --- | --- | --- | --- | --- | --- | --- |
| Marker | Dilution (μl) | Fluorophore | Laser | Company | Catalog Number | Clone |
| HLADR | 0.63 | BUV661 | UV11 | BD | 612980 | G46-6 |
| CXCR4 | 1.25 | BUV395 | UV2 | BD | 563924 | 12G5 |
| CD169 | 0.63 | BUV496 | UV7 | BD | 750364 | 7-239 |
| CD86 | 1 | BUV737 | UV14 | BD | 612785 | 2331 (FUN-1) |
| CD115 (CSFR1) | 5 | SuperBright436 | V2 | Thermo | 62-1159-41 | 12-3A3-1B10 |
| CD209 | 2 | BV421 | V1 | Biolegend | 330118 | 9E9A8 |
| CX3CR1 | 2.5 | SuperBright 702 | V13 | Thermo | 67-6099-42 | 2A9-1 |
| MERTK | 2.5 | BV711 | V14 | Biolegend | 367620 | 590H11G1E3 |
| CD127 | 2.5 | SuperBright 780 | V15 | Thermo | 78-1278-42 | RDR3 |
| CD123 | 0.63 | BV480 | V5 | BD | 566182 | 9F5 |
| CD11b | 0.31 | eFluor506 | V7 | Thermo | 69-0112-80 | M1/70 |
| CD18 | 2.5 | SuperBright 600 | V10 | Thermo | 63-0189-41 | 6.7 |
| CD56 | 1 | SuperBright 645 | V11 | Thermo | 64-0566-42 | WM-53 |
| CD45 | 0.5 | NFB610 | B6 | Thermo | H005T02B05 | 2D1 |
| CD16 | 1.25 | NFB555 | B3 | Thermo | H006T02B03 | 3G8 |
| CD14 | 2 | NFB660 | B7 | Thermo | H007T02B07 | MEM-15 |
| CD301 | 2.5 | AF488 | B2 | Thermo | MA5-23645 | 744812 |
| CD88 | 5 | PerCP-eFluor710 | B10 | Thermo | 46-0888-41 | S5/1 |
| CD89 | 2.5 | PerCP-Cy5.5 | B9 | Biolegend | 354110 | A59 |
| LOX1 (OLR1) | 1 | PE | YG1 | Biolegend | 358604 | 15C4 |
| CD204 | 5 | PE-Cy7 | YG9 | Thermo | 25-2045-41 | PSL204 |
| CD206 | 5 | PE-Cy5 | YG5 | Biolegend | 321108 | 19.2 |
| CD11c | 1.25 | PE-Cy5.5 | YG7 | Thermo | 35-0116-42 | 3.9 |
| CD54 (ICAM-1) | 0.625 | AF700 | R4 | Thermo | 56-0549-42 | HA58 |
| CD163 | 1 | AF647 | R2 | Thermo | 51-1637-42 | MAC 2-158 |
| VEGFR1 | 10 | APC | R1 | RnD | 49560 | FAB321A |
| CD31 | 0.63 | APC-eFluor780 | R7 | Thermo | 47-0319-41 | WM59 |

